# Language and the cerebellum: structural connectivity to the eloquent brain

**DOI:** 10.1101/2022.04.19.488812

**Authors:** Katie R. Jobson, Linda J. Hoffman, Athanasia Metoki, Haroon Popal, Anthony Steven Dick, Jamie Reilly, Ingrid R. Olson

## Abstract

Neurobiological models of receptive language have focused on the left-hemisphere perisylvian cortex with the assumption that the cerebellum supports peri-linguistic cognitive processes such as verbal working memory. The goal of this study was to identify language-sensitive regions of the cerebellum then map the structural connectivity profile of these regions. Functional imaging data and diffusion-weighted imaging data from the Human Connectome Project (HCP) were analyzed. We found that (a) working memory, motor activity, and language comprehension activated partially overlapping but mostly unique subregions of the cerebellum; (b) the linguistic portion of the cerebello-thalamo-cortical circuit was more extensive than the linguistic portion of the cortico-ponto-cerebellar tract; (c) there was a frontal-lobe bias in the connectivity from the cerebellum to the cerebrum; (d) there was some degree of specificity; and (e) for some cerebellar tracts, individual differences in picture identification ability covaried with fractional anisotropy metrics. These findings yield insights into the structural connectivity of the cerebellum as relates to the uniquely human process of language comprehension.

## Introduction

One of the most controversial issues in cognitive neuroscience today is whether the cerebellum deserves a seat at the table of the core language network. Neuromotor control of speech, which is important for both comprehension and production of language (Fischer & Zwaan, 2008), can be affected by cerebellar damage, with ataxic dysarthria being a common outcome of cerebellar damage (Spencer & Slocomb, 2007). However, there is growing evidence suggesting the cerebellum’s involvement in language goes beyond its well-established role in the motor aspects of speech and comprehension (Mariën et al., 2014). The most compelling findings come from studies of individuals with damage to the posterior cerebellum and a disorder known as cerebellar cognitive affective syndrome. Individuals with this disorder have disruptions in many cognitive functions, including features of language (Schmahmann & Sherman, 1998). Prior work has found that individuals with this disorder often produce agrammatic language output (De Smet et al., 2007; Mariën et al., 2000; Silveri et al., 1994), with concurrent deficits in sentence comprehension (De Smet et al., 2007; Murdoch & Whelan, 2007; Silveri, 2021), anomia as defined by deficits in word generation (Mariën et al., 2000; De Smet et al., 2007) and diminished verbal fluency (Hoche et al., 2018; Silveri, 2021). Although linguistic and motoric deficits are often co-morbid in acquired cerebellar injuries, these two domains are also dissociable (Ahmadian et al., 2019). The cerebellum has, however, proven to be an enigmatic lesion model. For example, people with acquired cerebellar injuries tend to experience rapid recovery of speech and language functions, with frank symptoms only apparent during the acute phase in adults (Fabbro et al., 2004).

Neuroimaging studies have supplemented and extended patient-based case studies. Recent studies have reported that language comprehension activates Lobule IV, Crus I & II and Lobule IX in the posterior cerebellum (Geva et al., 202; Vias & Dick, 2017), with a rightward cerebellar bias, due to the fact that cortico-cerebellar-cortical connections are crossed (Bostan, Drum & Strick, 2013). What is the computational role of this region in language? Several hypotheses have been offered. One idea is that the cerebellum modulates timing and sequencing of language production and language perception (Fiez, 2016; Ivry & Keele, 1989; Mariën et al., 2014; Mariën & Borgatti, 2018; Molinari et al. 1997; Molinari, 2015; Salman, 2002; Schwatze and Kotz, 2016). Another hypothesis is that the cerebellum is involved in automatizing a range of behaviors, including language (Doyon et al., 1998; Fiez, 2016; Ramnani, 2014; Vicari et al., 2018; Yang et al., 2014). A final hypothesis is that the cerebellum’s role in language is related to the more general role of the cerebellum in verbal working memory (Chen & Desmond, 2005; Desmond et al., 1997; Marvel & Desmond, 2010; Peterburs et al., 2021).

The comprehensive mapping of cortico-cerebellar connectivity can provide a framework upon which to understand function. More specifically, rather than understanding the role of the cerebellum in language more broadly, the mapping of the pathways that connect language-specific regions in the cerebellum and cerebral language regions may show some specificity. If this connectivity can be shown to be functionally specialized for particular linguistic sub-domains, it may show more specifically how the cerebellum contributes to each linguistic sub-domain. For over one hundred years it has been recognized that certain white matter tracts play a key role in language (Dejerine & Dejerine-Klumpke, 1895; Dejerine & Dejerine-Klumpke, 1901; Miraillé, 1896). Modern diffusion-weighted imaging (DWI) methods have allowed investigators to identify structural networks in the cerebrum essential for language (Dick & Tremblay, 2012; Duffau, 2015; Middlebrooks et al., 2017; Krestel et al., 2013; Smits, Jiskoot, & Papma, 2014). Whether cerebral language networks are structurally connected with portions of the cerebellum activated in language tasks is not known. Although functional connectivity studies have shown correlated activity between linguistically-sensitive cortex and regions of the posterior cerebellum (reviewed in Vias & Dick, 2017) this method is far removed from “ground truth” evidence provided by post-mortem tract tracing. Indeed, findings from functional connectivity often disagree with findings from structural connectivity (for an example, see Metoki et al., 2021).

Findings from tract-tracing have shown that cerebellar connectivity to and from the cerebrum is uniquely defined by two major pathways. The cortico-ponto-cerebellar pathway projects from the cerebral hemispheres with decussation at the pons terminating in the contralateral cerebellar cortex. In contrast, the cerebello-thalamo-cortical pathway begins in the cerebellum then crosses over to synapse on the contralateral thalamus, continuing on to different regions of the cerebral cortex. The dentate nucleus of the cerebellum retains a topographically ordered pattern of connectivity (Dum & Strick, 2003; Steele et al., 2016; Palesi et al., 2021). It is presumed that this connectivity remains orderly throughout the cerebello-thalamo-cortical and cortico-ponto-cerebellar pathway tracts.

The goal of this study is to use DWI to help disentangle the role of the cerebellum in language comprehension. DWI provides information about what brain regions are communicating with each other via a mathematical model of axonal connectivity. Information about axonal connectivity can be used to determine which facets of language are mediated by the cerebellum.

We used a tractography pipeline that we previously developed for studying the role of the cerebellum in theory of mind (Metoki et al., 2021). We applied this pipeline to language ROIs in the cerebellum and cerebrum. Cerebellar language ROIs were derived from the sentence comprehension task in the HCP task-fMRI data set (Binder et al., 2011). Cerebral ROIs were chosen a priori based on the sentence comprehension literature (Barch et al., 2013; Binder et al., 2011; Booth et al., 2007; Fengler et al., 2016; Friederici et al., 2003; Keller et al., 2001; Kieren & Buckner, 2009; Rogalsky et al., 2008). We focused on language comprehension, rather than overt language production, with the goal of reducing the potential impact of motor processing. Because language tends to be strongly left lateralized in the cerebrum (Frost et al., 1999; Mariën et al., 2001; Takaya et al., 2015) and the cerebellum has crossed structural connectivity with the cerebrum (Gonzalo-Ruiz, & Leichnetz, 1990; Ito, et al., 1986; Kelly & Strick, 2003), we performed lateralized tractography between language-sensitive ROIs in the right cerebellum to language-sensitive ROIs in the left cerebrum. Based on prior DWI work (Metoki et al., 2021) as well as monkey histology work (Glickstein, May, & Mercier, 1985; Schmahmann, 1996) we predicted that there would be significantly more fibers to linguistically-sensitive frontal cerebral ROIs, like the left inferior frontal gyrus, as compared to linguistically-sensitive ROIs in the temporal lobe.

## Materials and Methods

### Dataset and Participants

All data used in this study are part of the HCP dataset, specifically the WU-Minn HCP Consortium S900 Release (WU-Minn HCP Consortium, 2015). This dataset is publicly available and accessible at https://www.humanconnectome.org. Only participants that completed all imaging sessions of interest (T1/T2, task fMRI (tfMRI)), and DWI scans were included in this study. We restricted our population to only right-handed subjects using the Edinburgh Handedness questionnaire (Oldfield 1971). We chose a random subset of 100 participants (50 females, Mean age = 27.89 years, SD = 3.9 years), as using a large sample size incurs a computational cost, given that probabilistic tractography is mathematically intensive. Unless otherwise stated, all significant results reported in this study were corrected for multiple comparisons using the false discovery rate correction (FDR; Benjamini and Hochberg, 1995).

### Overview of HCP behavioral protocol

While known for its neuroimaging data, the HCP protocol also includes several behavioral assessments conducted outside of the scanner. The NIH Toolbox for Assessment of Neurological and Behavioral function was of most interest to us, in particular their tasks related to language comprehension. The NIH Toolbox Picture Vocabulary Test, a measure of receptive language, was our dependent behavioral measure. This particular version of the task was adaptive, which allowed for more variation in this data (for additional detail see Gershon et al. (2014)). Note that fMRI tasks are generally designed to produce strong BOLD signals but often have very little variance in the resultant behavioral outcomes. The HCP behavioral tasks used outside of the scanner elicited higher variance in the behavioral outcomes, making them potentially more sensitive for analyses of individual differences.

### Overview of HCP fMRI protocol

A detailed description of the HCP data acquisition and preprocessing pipelines can be found elsewhere (Van Essen et al., 2012; Barch et al., 2013; Glasser et al., 2013; Smith et al., 2013). Briefly, the HCP protocol includes acquisition of structural MRI, task-state fMRI, diffusion MRI, and extensive behavioral testing. The imaging data used in this article are the “minimally preprocessed” subjects included in the WU-Minn HCP Consortium S900 Release (WU-Minn HCP Consortium, 2015). This includes standard preprocessing using TOPUP, EDDY and BEDPOSTX. Details of imaging protocols, preprocessing pipelines, and in-scanner task protocols can be found in the Supplementary Material.

Task-state fMRI encompasses seven major domains; three of which were used in this study: (1) working memory/cognitive control systems; (2) motor (visual, motion, somatosensory, and motor systems); and (3) language comprehension. The main task of interest was the language task (Binder et al., 2011). Participants in the language task listened to stories adapted from Aesop’s fables (www.aesopfables.com). Sentences were read aloud by a text-to-speech program to participants in the scanner. After listening to the stories, the participants were presented with a two-answer forced choice question about the contents of the story. The question was meant to probe understanding about the theme of the story, thus evoking activations related to comprehension (Binder et al., 2011). Participants selected one of the two answers by pushing a button. Specific details about the contents of the stories such as number of events, number of actors, mean sentence length, and duration are described in detail by Binder and colleagues (2011). The accompanying control task involved participants doing math problems. The design of this task was the same as the language task, with the same text-to-speech method used to present the stimuli. Rather than listening to a story, participants were read math problems aloud (“six times two equals…”). Participants were presented with a mathematical problem and were asked a two-answer forced choice question. The difficulty of the math task was increased after six correct responses and decreased in difficulty after one incorrect response. Descriptions of the working memory and motor tasks, used in the overlap and control analyses, are further described in the Supplementary Material.

The results of the language fMRI task task in the scanner did not yield enough variance to examine its potential relationship with our white matter pathways. Instead, we included participants’ performance on the NIH Toolbox Picture Vocabulary Test as our dependent behavioral measure.

### Regions of Interest (ROIs)

We used two sets of ROIs for our analyses: one set in the cerebrum and one in the cerebellum. Because language is strongly lateralized, we limited our ROIs to the left cerebral hemisphere and to the right cerebellum. The set of cerebral ROIs were drawn from prior work on language. The cerebral ROIs included the following: angular gyrus (ANG; Fengler et al., 2016; Keller et al, 2001; Van Ettinger-Veenstra et al., 2016), dorsolateral prefrontal cortex (DLPFC; Keller et al., 2001; Kieren & Buckner, 2009), superior temporal gyrus (STG; Barch et al., 2013; Binder et al., 2011; Booth et al., 2007; Fengler et al., 2016; Friederici et al., 2003; Keller et al, 2001; Turken & Dronkers, 2011), middle temporal gyrus (MTG; Binder et al., 2011; Keller et al., 2001; Turken & Dronkers, 2011; Van Ettinger-Veenstra et al., 2016), inferior temporal gyrus (ITG; Ikuta et al., 2006), temporal pole (TP; Barch et al., 2013; Binder et al., 2011), inferior frontal gyrus/Broca’s (IFG; Barch et al., 2013; Binder et al., 2011; Booth et al., 2007; Van Ettinger-Veenstra et al., 2016) and posterior superior portion of the temporal lobe (PST; Just et al., 1996; Mesulam et al., 2015). This area is often considered to be synonymous with Wernicke’s Area; however, because there is a lack of consensus regarding its location in the field (Tremblay & Dick, 2016), throughout this paper we will refer to this area as the posterior superior temporal lobe (PST). Although other brain areas also support language, we focused on regions that have been consistently implicated in sentence comprehension. Coordinates for cerebral ROIs were taken from Neurosynth (https://neurosynth.org/) by searching for the name of the region under ‘terms’ and taking the voxel with the highest z-score using a cluster analysis. Using the term ‘superior posterior’ produced an activation map closest to the agreed-upon anatomical location of Wernicke’s Area (Tremblay & Dick, 2016). Unfortunately, the highest z-scored voxel produced under the term ‘posterior superior’ was too posterior for the aims of this study. Instead, we took the highest z-scored voxel that was also close enough to Heschl’s gyrus as justification for using it to represent the PST. See Table A in the Supplementary Material for exact MNI coordinates and accompanying z-scores for each ROI. Subsequently, each coordinate was then transformed into a 6mm sphere using FSL software, to be used in the analysis.

For the ROIs in the cerebellum, we used a different apporach. Although several studies have attempted to functionally map the cerebellar cortex (Buckner et al., 2011; Diedrichsen & Zotow, 2015; Guell et al., 2018a; 2018b; King et al., 2019; Krienen & Buckner, 2009; Marek et al., 2018; Riedel et al., 2015), there is no consensus about functional boundaries within the cerebellum. Hence, it was impossible to employ the same approach as we did with the predefined cerebral ROIs. Instead, we used a data-driven approach. Following the method used by Guell et al. (2018a), we transformed FSL’s level 2 individual cope files (results of within-subject fixed-effects grayordinate-based analyses which generate output files that index mean effects for an individual subject averaged across the 2 scan runs for a task) into Cohen’s d group maps by first transforming the grayordinate.dscalar.nii files to NIfTI. We then used FSL commands fslselectvols to extract the contrast of interest “story > math” for each individual, and fslmerge, fslmaths -Tmean, -Tstd, and -div to merge the individual contrast images, extract the mean, and the standard deviation, and divide the two, ultimately getting group Cohen’s d maps for the contrasts “story > math” (language), “2-back > 0-back” (working memory), and “average” (motor) based on a larger sample of 671 subjects. The HCP S900 Release provides level 3 group z-maps, but Cohen’s d maps made it possible to observe the effect size of each task contrast rather than the significance of the BOLD signal change. A sample of 671 subjects ensures that a d value higher than 0.5 (Cohen 1988) will be statistically significant even after correction for multiple comparisons (d = z/sqrt(n), d > 0.5 we have z > 12.95 for N = 671; analysis of 17,853 cerebellar voxels would require p < 0.000028 after Bonferroni correction, and p < 0.000028 is equivalent to z > 4.026). Accordingly, we used FSL’s cluster tool, the Cohen’s d maps, and a threshold of 0.5 to extract clusters of activation for each task and local maxima within each cluster. After using a whole cerebellar mask to retain only the clusters and local maxima within the right cerebellum, clusters smaller than 100 mm^3^ were further removed to omit very small clusters that were considered to be noninformative and would make a comprehensive description of the results too extensive (see Figure 1 for a visualization of the three functional tasks). The coordinates of the remainder local maxima from the “story > math” (language) contrast within the cerebellum were used to create two group cerebellar ROIs (spheres, 6 mm radius) in the right hemisphere. This resulted in two language cerebellar ROIs: right Crus I and right Lobule IX. The same method was used to extract the cerebellar motor and working memory cerebellar ROIs, which were used in overlap and control analyses. Seven working memory ROIs were created, also in the right cerebellar hemisphere. Four of the seven cerebellar working memory ROIs overlapped with each other, so they were removed from the analysis, leaving three working memory ROIs in the cerebellum. The remaining cerebellar working memory ROIs were primarily located in Lobule IV, Crus I, and Lobule VIIb.

**Figure 1.**
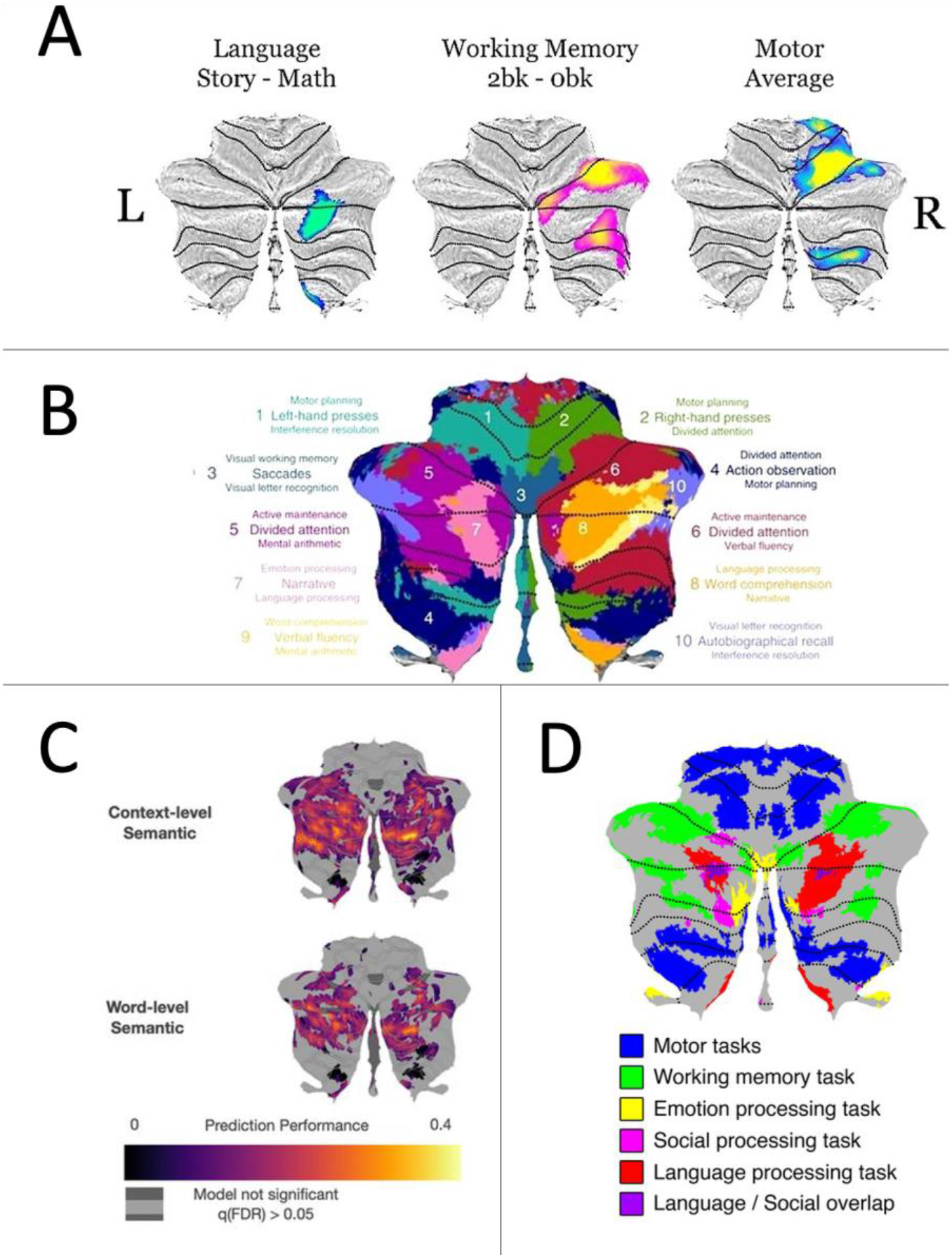
A) Functional activations to three tasks (language, working memory, and motor) displayed on a flatmap of the cerebellum. Only activations within the right cerebellar hemisphere are displayed. B) A figure from King et al (2019) depicting Cb activations to a range of cognitive and motor tasks. Note that there is overlap between their language activations (labeled primarily as numbers 7 and 8) and our language activations. C) A figure from LeBel et al (2021) investigating sentence- and word-level semantic language processing. D) A figure from Guell et al. (2018), displaying Cb activations to a range of cognitive and motor tasks from the HCP dataset. Note the similar activation patterns, as Figures 1A and 1D come from the same dataset.

### Cluster Overlap and Euclidean distances

Given that the HCP dataset uses FNIRT registration to the MNI template, we calculated the percentage of overlap of each cerebellar cluster by using Diedrichsen’s FNIRT MNI maximum probability map (Diedrichsen et al. 2009). Previous studies have already explored the overlap of several other tasks included in the HCP dataset (Metoki et al., 2021). As a result, we only included three tasks in this analysis: language, working memory, and motor. We used FSL’s atlasq tool to determine the percent overlap of each cerebellar cluster in the cerebellar lobes, hence determining the primary location of each cluster. The Sørensen–Dice coefficient, which is a statistic measuring the similarity of two samples (Dice, 1945; Sørensen, 1948), was then used to calculate the percentage of overlap between the functional clusters generated from all tasks and determine their similarity, and Euclidean distances were calculated to estimate the distances of local maxima within and between clusters. At the individual level, we thresholded *Z*•-scored *β*-weights of each subject’s activation map for each task contrast to > 0 to retain only increased activation during the tasks and then ran Wilcoxon signed-rank tests between each task pair to examine whether there was a statistical difference between them.

### Diffusion Analyses

All diffusion analyses were completed on Temple University’s High-Performance Computing Cluster, OwlsNest. Probabilistic tractography analyses were performed using FSL’s probtrackx2 (probabilistic tracking with crossing fibres, version 6.0.2) (Behrens et al., 2003, 2007) in each subject’s native space. Due to HCP preprocessing steps, this native space we refer to is the subjects’ T1w space. After tractography, results were then transformed to MNI standard space using transformation matrices (see Supplemental Material for more details). An ROI-to-ROI approach was used with cerebral and cerebellar ROIs used as seeds and targets to reconstruct each subject’s cerebello-cerebral white matter connections. Fiber tracking was initialized in both directions separately (from seed to target and vice versa) and 5,000 streamlines were drawn from each voxel in each ROI. Tract length correction was also used, as cerebellar tracts are lengthy due to their polysynaptic nature, and length of a tract can introduce more false positives to the data (Jones, 2010). Tractographies were performed to delineate the cerebello-thalamo-cortical pathway, which projects from the cerebellar cortex to the deep cerebellar nuclei then crosses over to synapse on the contralateral thalamus, continuing on to different regions of the cerebral cortex (Middleton & Strick, 1997; Schmahmann & Pandya, 1997; Palesi et al., 2015) and the cortico-ponto-cerebellar pathway, which projects from the cerebral hemispheres to the pons, then to the contralateral cerebellar cortex (Ramnani, 2006; Palesi et al., 2017). For the cerebello-thalamo-cortical tractographies, a binarized mask of the superior cerebellar peduncle in MNI space from the Johns Hopkins University ICBM-DTI-81 white-matter labels atlas (Mori, Wakana, Van Zijl, & Nagae-Poetscher, 2005; Wakana et al., 2007; Hua et al., 2018) and left thalamus in MNI space from the Harvard-Oxford subcortical atlas (Frazier et al., 2005; Makris et al., 2006; Desikan et al., 2006; Goldstein et al., 2007) were used as waypoints respectively. The binarized contralateral cerebellar and cerebral hemispheres, lobes of non-interest, and the opposing cerebellar peduncle were set as exclusion masks. All cerebral masks were created using the Harvard-Oxford cortical atlas (Frazier et al., 2005; Makris et al., 2006; Desikan et al., 2006; Goldstein et al., 2007). For example, cerebello-thalamo-cortical tractography between the right cerebellum and left DLPFC included the right superior cerebellar peduncle and left thalamus as waypoints. The exclusion mask comprised of the left cerebellar hemisphere, right cerebral hemisphere, middle cerebellar peduncle, precentral gyrus, as well as the temporal, parietal and occipital lobes. For the cortico-ponto-cerebellar pathway tractographies, a binarized mask of the middle cerebellar peduncle in MNI space from the same atlas was used as a waypoint. The contralateral cerebellar and cerebral hemispheres, right superior cerebellar peduncle and lobes of non-interest were used as exclusion masks. For example, cortico-ponto-cerebellar pathway tractography between a left temporal lobe ROI and right cerebellar ROI would entail the middle cerebellar peduncle waypoint mask. The exclusion mask included the left cerebellar hemisphere, right cerebral hemisphere, right superior cerebellar peduncle, precentral gyrus, and frontal, parietal and occipital lobes. Exact regions included in each exclusion mask can be found in Supplementary Table B. The pons was not selected as an inclusion mask due to its absence in any standardized atlases. Despite not having the pons as an orthogonal waypoint in the cortico-ponto-cerebellar pathway tract, we did include the thalamus in the cerebello-thalamo-cortical reconstruction to follow previous literature (Palesi et al., 2015) and to ensure our tractography was as anatomically similar to the ground truth as possible. Our exclusion masks were comprehensive, as we had certain expectations as to where the tract would be traveling and terminating (Schilling et al., 2020). For cerebral ROIs in the frontal lobe, we knew that we were not interested in fibers that extended into other regions such as the parietal or temporal lobes, therefore we incorporated them in the exclusion mask for that ROI. We also excluded the motor cortex with the knowledge that there are tracts from the cerebellum to the motor cortex. Exclusion of the motor cortex ensured our results were not due to connections between cerebral motor regions and our ROIs in the cerebellum.

Two metrics were extracted to be used as dependent measures in our analyses. Volume was extracted from the streamline density map using FSL’s fslstats and normalized using intracranial volume (Voevodskaya et al., 2014). Intracranial volume was calculated using Malone et al’s (2015) method. This involves isolating the skull-stripped T1-weighted image provided by HCP in the same native space as the diffusion data and segmenting the brain using SPM12 into gray matter, white matter, and CSF (in liters). The summed volumes of these tissue types yields the whole-brain intracranial volume, which when transformed into mm^3^ yields the metric for probabilistic tractography reported by FSL. Additionally, fractional anisotropy (FA) was extracted from respective cerebellar peduncles and included as a dependent variable. This was accomplished by taking whole-brain FA scalar data in native space (the result of the command dtifit, which is run on eddy-corrected data) and using each of the peduncles to mask the data. This produced FA maps exclusively within each peduncle. That data was then transformed into standard space, which allowed us to extract microstructural information of each tract within each peduncle. We chose to extract microstructural indices from only the peduncles following previous literature, as the peduncles are generally a good point in the tract to evaluate white matter cohesion (Jossinger et al., 2021; Taoka et al., 2007; Wang et al., 2003). Selecting only one region along the resulting tractography should reduce some of the noise that would be introduced at the termination of the tract, which is a concern for such lengthy fiber pathways. This also allowed us to isolate the contribution of the cerebellar pathways in behavior, without picking up on cerebral white matter pathways (such as the arcuate fasciculus).

## Results

### Functional domains in the Cerebellum

We first examined whether there was overlap between language, working memory and motor activations in the right cerebellum. We found that there was a significant difference in localizations for all three task pairs. Overall motor activations were localized to the anterior cerebellum (Lobules I-IV), with some activation in the posterior cerebellum (lobules VIIB and VIIIA). Overall language activations were localized to Crus I/II and lobule IX. Overall working memory activations were found in Crus I/II, Lobule VI, Lobule VIIb and Lobule VIIIa. The *β*-weights of each subject’s activation map that were previously extracted, we performed Wilcoxon signed-rank tests for each task pair to examine whether there was a statistical difference between them. The language and motor activations had no overlap, while the language and working memory functional activations had minimal overlap (0.005%). After conducting the analyses on the overall activations from these tasks, the peak activation was extracted from the group activation maps, and transformed into 6mm spheres in FSL to be used in subsequent analyses.

### Cerebellar Structural Connections: language-language

Next, we asked whether language-sensitive regions in the cerebellum, which are poorly understood and have received little attention, are structurally connected to language-sensitive areas in the cerebrum that have been studied for over a century. We ran probabilistic tractography to reconstruct the cerebello-thalamo-cortical and cortico-ponto-cerebellar pathway white matter pathways between ROIs in the right cerebellum and left cerebral cortex. Volume was extracted for the cerebello-thalamo-cortical and cortico-ponto-cerebellar pathways and FA (microstructure) was extracted for the superior (cerebello-thalamo-cortical tracts) or middle (cortico-ponto-cerebellar tract) peduncles.

The average volume for tracts (before correction for intracranial volume) ranged from 74,714.32mm to 240,083.92mm. There were significant differences between the cerebello-thalamo-cortical and cortico-ponto-cerebellar pathways with lower volume in the cortico-ponto-cerebellar pathways (see Table 2 & Figure 2; W = 8614, *p* < .001). For the cerebello-thalamo-cortical pathway, we examined whether there was a difference in the average volume between each cerebellar ROI seed (Crus I and Lobule IX). For this analysis, we intended to isolate the average volume when a tract was seeded in a cerebellar region to identify if one seed had a relatively greater volume of tracts being sent to the cerebrum. We accomplished this by taking each tract seeded in its respective cerebellar ROI to each of the eight cerebral targets (IFG, DLFPC, ANG, TP, STG, MTG, PST, ITG) and averaging the numbers. Using a paired Wilcoxon signed-rank test, we found no significant difference in tract volume based on cerebellar seed. Paired Wilcoxon signed-rank tests were then used to look at differences in volume based on target cerebral ROIs for cerebello-thalamo-cortical pathways. We found that tracts terminating in frontal lobe ROIs (IFG and DLPFC) had significantly higher volumes than those of any terminating in other lobe (see Figure 3A). There was no difference between the parietal and temporal lobe tract volumes, but we found some diversity amongst temporal lobe ROIs. We found a significant difference between MTG and PST (V = 1139, *p* = 0.037), ITG and PST (V = 880, *p* < 0.001), as well as between the TP and PST (V = 997, *p* < 0.001). There was also a significant difference between IFG and DLPFC volumes, with IFG having the greater volume between the two (V = 699, *p* < 0.001).

**Table 1.** The Sørensen–Dice coefficient (in percentages) between the three functional activations for the tasks of interest in the right cerebellar hemisphere. All three tasks showed very little overlap. Representation of the activations is depicted in Figure 1.

| **fMRI Task Activation** | **Language** | **Working Memory** | **Motor** |
| --- | --- | --- | --- |
| *Language* |  |  |  |
| *Working Memory* | 0.0051 |  |  |
| *Motor* | 0 | 0.1507 |  |

**Table 2.** A summary comparison of the volumes and standard deviations (SD) for the cortico-ponto-cerebellar and cerebello-thalamo-cortical connections to language-related cortical regions. Note that the cerebellar region of interest for all tracts is Crus I for the below delineated descriptives.

| Language ROI | CTC (seeded in Crus I) |  |  |  | CPC (seeded in Crus I) |  |  |  |
| --- | --- | --- | --- | --- | --- | --- | --- | --- |
|  | Uncorrected Vol. |  | ICV Corrected Vol. |  | Uncorrected Vol. |  | ICV Corrected Vol. |  |
|  | Mean | SD | Mean | SD | Mean | SD | Mean | SD |
| *STG* | 150664.72 | 55440.85 | 0.10 | 0.04 | 91803.12 | 26480.96 | 0.06 | 0.02 |
| *MTG* | 150863.84 | 55557.13 | 0.10 | 0.04 | 85077.28 | 30491.50 | 0.05 | 0.02 |
| *ITG* | 150506.24 | 55179.80 | 0.10 | 0.04 | 74621.92 | 27725.44 | 0.05 | 0.02 |
| *ANG* | 150000.88 | 54587.71 | 0.10 | 0.04 | 117644.72 | 30662.77 | 0.08 | 0.02 |
| *TP* | 150073.92 | 55120.73 | 0.10 | 0.04 | 123634.80 | 28211.70 | 0.09 | 0.02 |
| *IFG* | 226984.88 | 74864.76 | 0.16 | 0.06 | 175971.60 | 41002.20 | 0.08 | 0.03 |
| *DLPFC* | 228344.00 | 75418.98 | 0.16 | 0.06 | 122568.64 | 45010.97 | 0.12 | 0.03 |
| *PST* | 151325.60 | 55894.72 | 0.10 | 0.04 | 71870.72 | 29796.13 | 0.05 | 0.02 |

**Figure 2.**
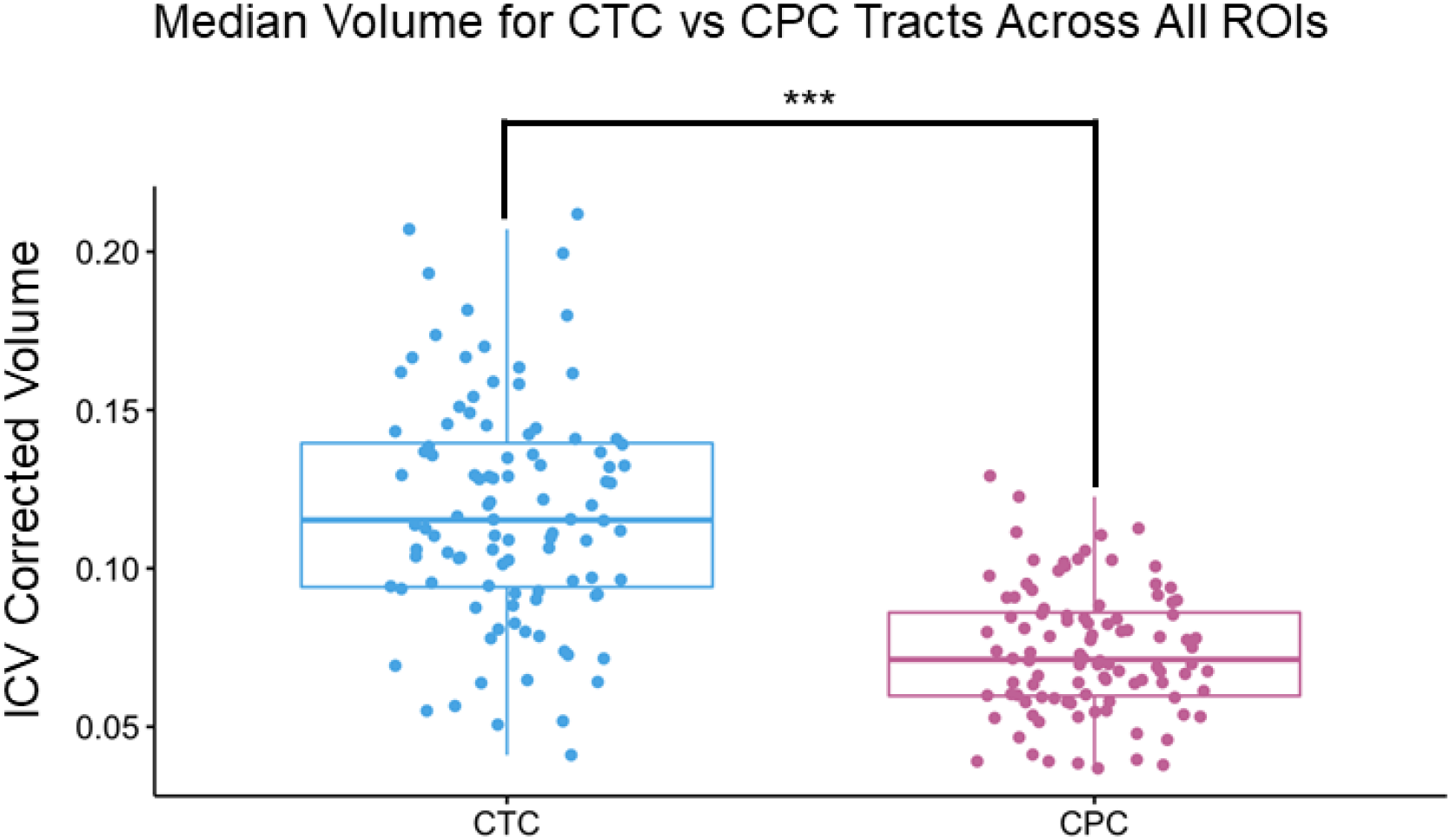
Median volume of the linguistic portion of the cerebello-thalamo-cortico (CTC) and cortico-ponto-cerebellar (CPC) tracts. Each pathway was created by averaging the volume of all tracts across all eight cerebral ROIs, resulting in a single average volume per tract for each participant. For example, the CTC seeded in Crus I had tracts calculated for each of the right cerebral ROIs. All eight of those tract volumes were averaged to create the CTC plot shown in this figure. Note that the size of the ROIs was identical in both analyses. For this particular example, pathways seeded/terminated in Crus I. This was done for illustration purposes. Each dot represents a single participant; *** indicates p < .001.

We found a different pattern of connection for the cortico-ponto-cerebellar pathway pathway (see Figure 3B). Again using paired Wilcoxon signed-rank tests, we found that there was still a statistically significant difference between the IFG and all other cerebral ROIs. Additionally, there were differences in volume in the temporal and parietal ROIs. Rather than the frontal ROIs being greater than the rest and all other tracts being equal to each other, we found that tracts from the angular gyrus, STG, and TP seeds had higher volumes than in the cerebello-thalamo-cortical tract. The overall pattern was IFG>DLPFC⩭ANG⩭TP>STG⩭MTG>WER⩭ITG (with ⩭ signaling statistical equivalence). Overall for both tracts, the tracts whose cerebral ROIs were IFG/Broca’s Area had the highest volume (significant at *p* < .001 in all pairwise comparisons - see Supplemental Table C for full list of comparisons).

**Figure 3.**
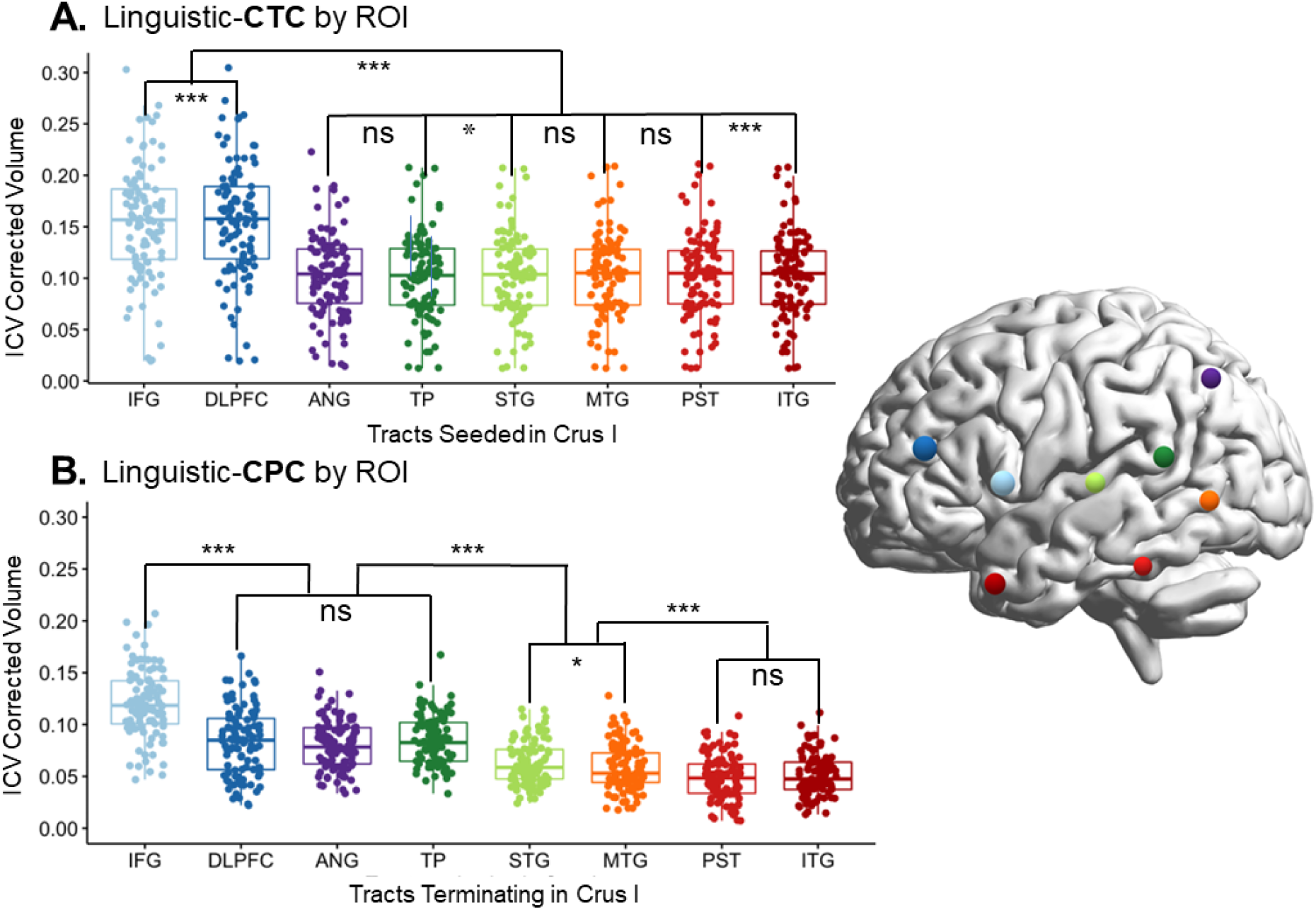
Volume comparison A. Between tracts beginning in right Crus I and ending in different left-lateralized cerebral language targets; and B. Between tracts projecting from cerebral language ROIs to the target in Crus I. CTC = cerebello-thalamo-cortical pathway; CPC = cortico-ponto-cerebellar pathway; IFG = inferior frontal gyrus/Broca’s; DLPFC = dorsolateral prefrontal cortex; ANG = angular gyrus; PST = posterior superior temporal lobe; STG = superior temporal gyrus; MTG = middle temporal gyrus; ITG = inferior temporal gyrus; TP = temporal pole. For this particular example, pathways seeded/terminated in Crus I. This was done for illustration purposes. For further information about each individual pairwise comparison, see Supplementary Table C; *** indicates p < .001, ** indicates p < 0.05, ns = non-significant.

### Cerebellar Structural Connections: language-working memory

To examine the specificity of these white matter connections, we ran probabilistic tractography between the cerebellar working memory ROIs and cerebral language ROIs using the exact same waypoints and exclusion masks as in the language-language analyses. The language-language connections were then compared to the working memory-language connections. We examined the white matter tracts projecting to and from IFG, as this was our highest volume target and seed. Results showed that the white matter pathways from the working memory cerebellar ROIs to the language cerebral ROIs had significantly lower volumes for tracts going to IFG in the cerebello-thalamo-cortical tracts (all comparisons p < .001; see Supplementary Table Table C for all p-values and effect sizes, and Table 3 for WM tract means). Interestingly, in the cortico-ponto-cerebellar pathway tracts, there was one exception: working memory ROI in right Lobule VI was not significantly different in ICV corrected volume than language ROI right Crus I (W = 2725, *p* = 0.667). These results suggest that there are specific structural connections between the linguistic cerebellar and cerebral areas, with the possibility that multiple regions of the cerebellum receive input from the same cerebral areas.

**Table 3.**
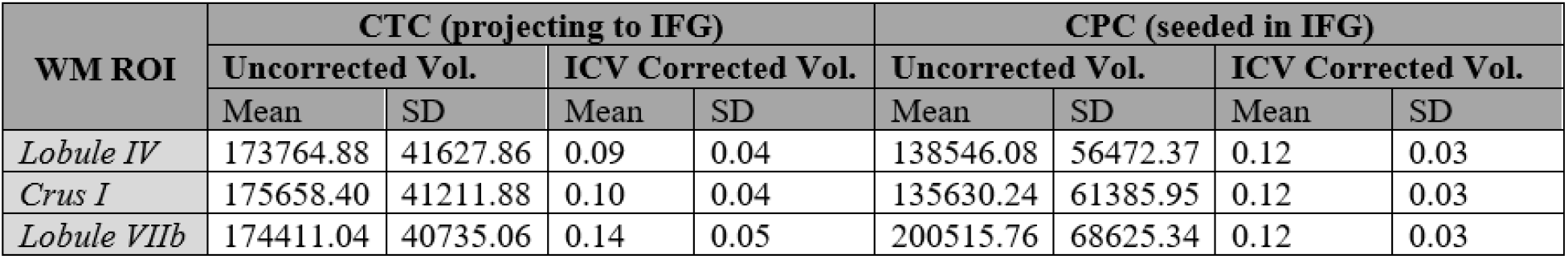
A summary comparison of the volumes and standard deviations (SD) for subsections of the cortico-ponto-cerebellar and cerebello-thalamo-cortical connecting to working memory (WM) regions. For the cortico-ponto-cerebellar projections to WM regions in the cerebellum are seeded in the IFG, while in the cerebello-thalamo-cortical working memory cerebellar regions are projecting to IFG.

### Cerebellar Microstructure: brain-behavior correlations

Next, we investigated the microstructural properties of each tract to determine if we could find a relation between white matter and behavior. We used Spearman’s rho to correlate FA extracted from the peduncles for each tract and the behavioral outcomes of the NIH Toolbox Picture Vocabulary Test from the HCP dataset. Spearman’s rho was used as some of our diffusion data from the cortico-ponto-cerebellar tract violated tests of normality (STG to Crus I: W = 0.964, *p* = 0.024; MTG to Crus I: W = 0.970, p = 0.057); STG to Lobule IX: W = 0.968, p = 0.043; MTG to Lobule IX: W = 0.977, p = 0.058). No significant correlations were found between microstructure of the cerebello-thalamo-cortical tract and picture naming ability. However, within the cortico-ponto-cerebellar tract there were several trending or significant correlations. For the tracts that had Crus I as the target, tracts seeded in: IFG (*r* = .23, *p* = 0.058), STG (*r* = .27, *p* = 0.020), and TP (*r* = .24, *p* = 0.041) were either trending or significant. For tracts with Lobule IX as the target, tracts seeded in: IFG (*r* = .24, *p* = 0.044), STG (*r* = .26, *p* = 0.024), and TP (*r* = .23, *p* = 0.050) were also significant.

### General Discussion

In this study we mapped structural pathways between language-sensitive regions in the cerebellum and language-sensitive regions in the cerebral cortex. Our first step was to identify regions of the cerebellum sensitive to verbal comprehension and then to examine overlap with motor and working memory task activations. The regions activated to the sentence comprehension contrast were localized to Crus I/Crus II, and Lobule IX (see Figure 1) of the posterior cerebellum. The location of these activations is consistent with prior work that was conducted on different participants and using different stimuli (King et al., 2019; Geva et al., 2021). It should be noted that the activation in Lobule IX has been a controversial addition to the linguistic cerebellum. However there is one piece of intriguing evidence linking it to receptive language: Geva and colleagues (2021) found that in their sample of six cerebellar lesion patients, damage to Lobule IX was the only lesion location that produced lasting deficits in sentence comprehension. Future research should examine this association more closely.

We found marginal overlap between the language and working memory tasks in Crus I, which is consistent with past literature (Marvel & Desmond, 2010). Τhis overlap could be attributed to the nature of the language task. Participants listened to fables and later made judgements about the fables. This required them to remember the gist of five to nine sentences, a task that invokes verbal working memory. The fact that these fables also involve social scenarios explains our previous finding that the language task has nearly 50% overlap with activations from the theory of mind task (Metoki et al., 2021). There is a sizable literature linking portions of the cerebellum to normal and abnormal social cognition as observed in autism spectrum disorder (Stoodley & Tsai, 2021; Van Overwalle et al., 2020). Whether language tasks that are less social and have lower working memory loads (e.g. single word processing) would show overlap with language activations in the cerebellum is unknown. In contrast, there was no overlap between the language comprehension task and the motor task. One explanation of the social and language overlap could be derived from the social sequencing hypothesis (Van Overwalle et al., 2019).This hypothesis states that the Cb is involved in the prediction of social interaction by forming internal models (a representation of the predicted world based on previous experiences) about how sequential social events should unfold. The Cb uses associative information to identify what may come next in a sequence of events, such as another person’s reaction or response. This relates to language because language is inherently social. Not only do we use it to relay information to another individuals, but the content of what we are relaying is often social. One study reported stronger activations in the Cb to social as compared to nonsocial sentences (Pu et al., 2020), adding credence to this idea. While the focus of our study was to parse receptive language and working memory, future work will need to be done to parse language and social cognition.

The observed overlap between language and working memory led us to conduct a distance analysis. This showed that each cluster had distinct local maxima. We also analyzed the *β*-weights for each task at the individual level and found that all task pairs have a significantly different localization of activation. Overall, our results provide support for the hypothesis that the cerebellum contains domain-specific mapping of cognitive functions including language comprehension.

If these cerebellar regions truly play a role in language, they should be structurally connected to regions involved in language in the cerebrum. Histology methods in macaques have revealed an extensive network of fiber paths between the anterior cerebellum and nearly all non-motor regions of the frontal lobe (Clower et al., 2005; Dum & Strick, 2003; Ito, 1984; Kelly & Strick, 2003; Leiner et al., 1993; Middleton & Strick 1997; Schmahmann & Pandya, 1997). In regards to the parietal lobe, gold-standard histology studies have found connections between the cerebellum and BA 5 and 7 in the macaque parietal lobe (Clower et al., 2005; Glickstein et al., 1985). Other histology studies in macaques have reported the existence of structural connections between the superior temporal lobe (e.g. the length of the superior temporal gyrus (excepting A1), as well as the depths of the superior temporal sulcus), with the cerebellum (Schmahmann & Pandya, 1991). However there are little to no connections between most of the middle temporal gyrus and inferior temporal gyrus and the cerebellum (Schmahmann & Pandya, 1991). Thus, histological findings in macaques predict that we should find strong structural connectivity between the cerebellum and language ROIs in the frontal lobe, and superior temporal lobe, but weaker structural connections between inferior parietal lobe and inferior temporal lobe ROIs and the cerebellum. This prediction must be tempered by the fact that language is a uniquely human trait, and changes have occurred in the human temporal and inferior parietal lobes through the evolution of language.

The results of our probabilistic tractography analysis partially confirmed the above predictions. First, we found that the volume of the language-specific cerebello-thalamo-cortical pathway was greater than the volume of the language-specific cortico-ponto-cerebellar pathway. This was not entirely unexpected as previous research found this bias when looking at the social portions of the cerebello-thalamo-cortical and cortico-ponto-cerebellar tracts (Metoki et al., 2021). Interpretation of this bias awaits further investigation.

Second, we found that in the cerebello-thalamo-cortical pathway, frontal lobe ROIs captured the majority of the projections from the cerebellum (IFG followed by the DLPFC) (see Figures 3 & 4). There was some elevated connectivity to the PST compared to other temporal lobe ROIs, but nothing compared to the volume of the pathways projecting to the frontal lobe, as was expected based on previous literature (Palesi et al., 2015).

**Figure 4.**
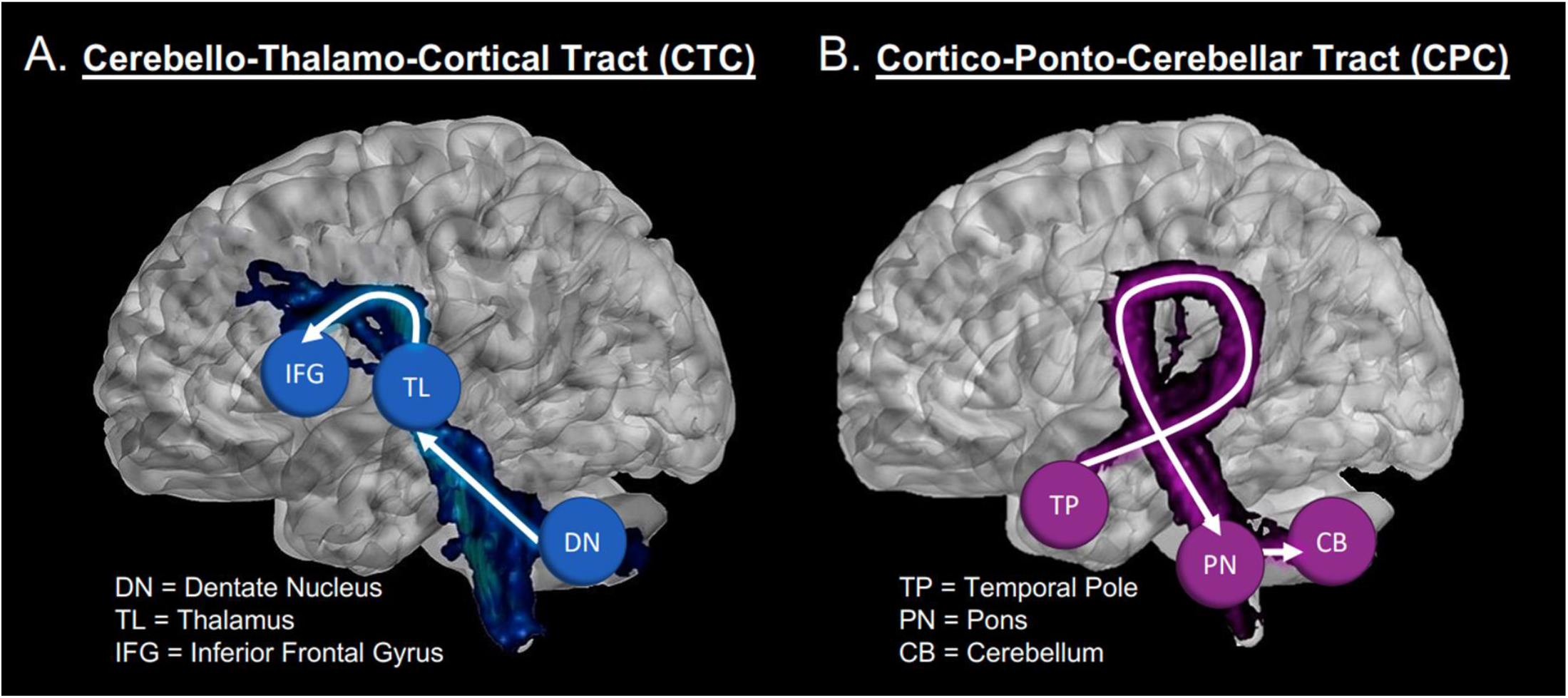
Visualization of: (A) The cerebello-thalamo-cortical tract running from Crus I of the cerebellum to inferior frontal gyrus (blue sphere); and (B) The cortico-ponto-cerebellar tract, running from the temporal pole (red sphere) to cerebellar Crus I.

**Figure 5.**
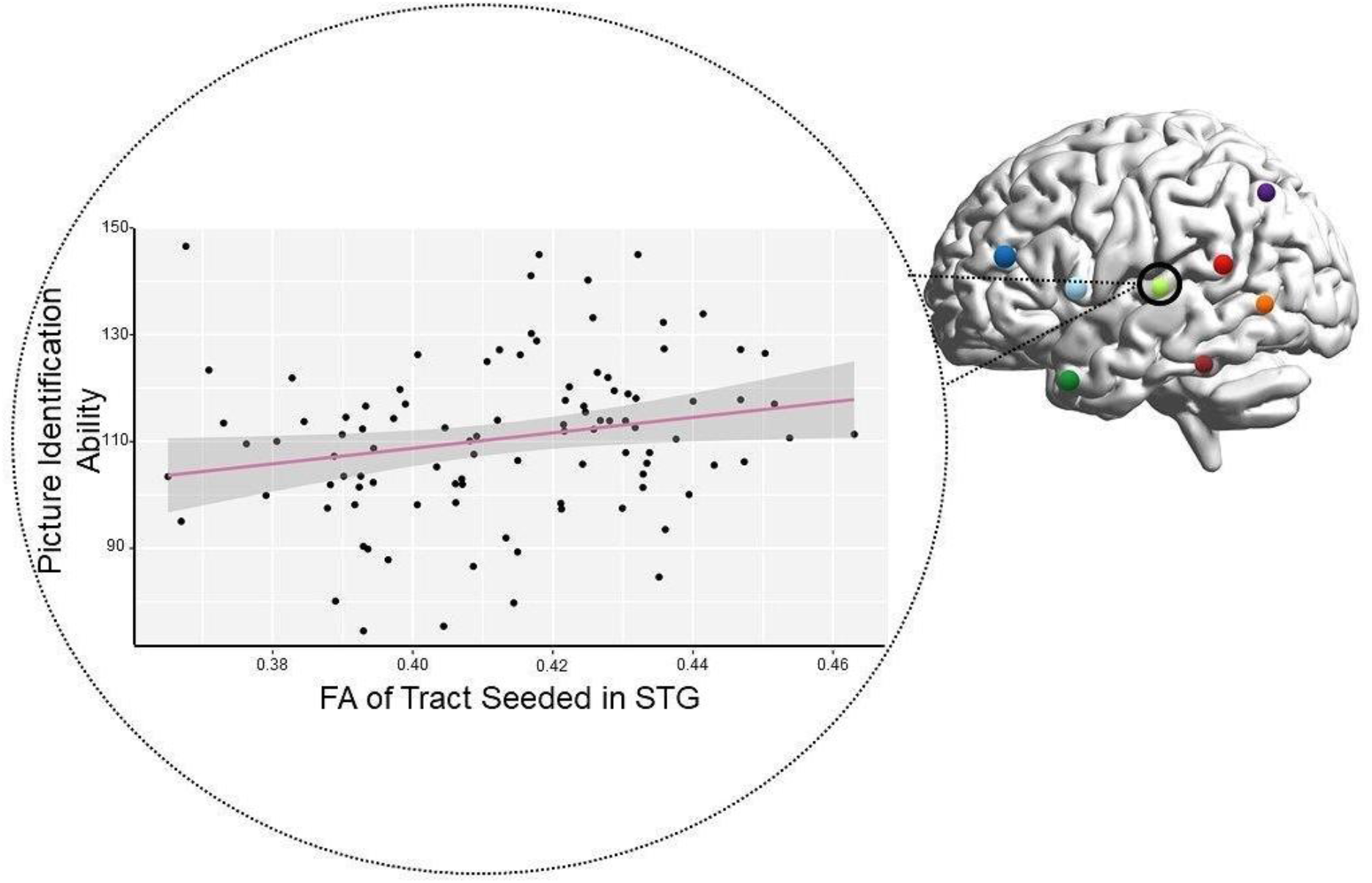
Correlation between microstructure (FA) of the STG-portion of the cortico-ponto-cerebellar tract plotted on the x-axis and single-word comprehension on the y-axis. Each dot represents a single participant.

However, the cortico-ponto-cerebellar pathway showed a different, more complex pattern of connectivity. While IFG still held the highest number of connections to the cerebellum, DLPFC, ANG, STG, and TP had elevated connectivity compared to other temporal lobe ROIs. The ANG and TP captured on average the same amount of volume as the DLPFC. The results involving the ANG and STG are not surprising, based on our review of the literature (Schmahmann & Pandya, 1991; Ramnani, 2012). However, finding elevated volume in the projections coming from the TP were unexpected. There is evidence from histology in macaques that the TP section of the superior temporal sulcus (TPO1) sends projections to the pons (Schmahmann, 1996), and likely onwards into the cerebellum.

These results provide new insight on the “closed-loop” nature of cerebellum-to-cerebrum connections. Computational models describing the cerebellum’s functional contribution to motor processes have described a closed loop system in which motor regions in the cerebrum are connected to specific regions within the cerebellum, which in turn are connected to the same regions back in the cerebrum (Ito, 2008; Kawato & Gomi, 1992; Wolpert et al., 1998). Evidence of closed-loop cerebello-cerebral connections that would be critical for non-motor processes have also been found (Palesi et al., 2017; Salmi et al., 2010; Strick et al., 2009), including closed-loops between the posterior cerebellum and the DLPFC (Kelly & Strick, 2003). However, as noted by Moberget and Ivry (2016), “… a close examination of the primate data suggests that [closed loops] may not always be the case.” The asymmetry between the cerebello-thalamo-cortical and cortico-ponto-cerebellar pathway in our functionally defined ROIs suggest that multiple regions across the cerebral cortex may be providing input into a specific region of the cerebellum, which then outputs to a subset of those input regions. This finding highlights that the closed-loop nature of cerebellar connectivity may be more in line with a network modulation function - a many to one - rather than a single region to region, or one to one, modulation.

Individual regions of the cerebellum would be well-suited to receive inputs from a diverse set of brain regions, as cerebellum “microcomplexes”, thought to be the computational building blocks of the cerebellum, include an initial layer of cells that are well-suited for high-dimensional inputs (Hull, 2020; Raymond & Medina, 2018).

To date, only a small number of studies have examined the cerebello-thalamo-cortical and cortico-ponto-cerebellar pathways *in vivo* (Karavasilis et al., 2019; Keser et al., 2015; Jissendi et al., 2008; Sokolov et al., 2014). This is likely due to the technical challenges present in doing tractography along a long, polysynaptic pathway that has crossovers and sharp turns. Thus most of the existing literature has simply looked at white matter within the cerebellum (e.g. the peduncles) rather than between the cerebellum and cerebrum. Our study is unique in that we carefully measured the linguistically-relevant portion of the ouroboros loop in a large sample with excellent data quality.

Previous studies that reconstructed cerebello-thalamo-cortical and cortico-ponto-cerebellar pathways relied solely upon diffusion *tensor* imaging, which has had notable issues with crossing fibers (Lee et al., 2015). Crossing fibers are in abundance in the brainstem and within these tracts of interest. Our tractography has improved upon this with the use of the ball-and-stick model, which is better at resolving crossing fibers (Behrens et al., 2007). We modeled our methods after Paelsi et al. (2017) who found evidence of projections of the cortico-ponto-cerebellar pathway originating from the temporal lobe with constrained spherical deconvolution. This method also has a better ability to solve for crossing fibers (Daducci et al., 2013; Tournier, Calamante & Connelly, 2012). Their analysis of all projections from association cortices showed that the temporal lobe sends projects to the cerebellum via the cortico-ponto-cerebellar pathway, even though it may not receive projections from the cerebello-thalamo-cortical. Schmahmann and Pandya (1991) also observed projections from the superior temporal sulcus to the cortico-ponto-cerebellar pathway in rhesus monkeys, so the results are not completely unfounded. It is possible these findings may be the result of the software and not true projections found in the human brain, but converging evidence across multiple methodologies may suggest some shared truth.

In an exploratory analysis of DWI microstructure data, we found that individual differences in FA values in the cortico-ponto-cerebellar pathway correlated with receptive vocabulary ability. In other words, individuals with higher FA in the middle cerebellar peduncle of the cortico-ponto-cerebellar tract had better ability to identify a picture that matched a word with which they were presented. Note that all tracts projecting from regions of the temporal lobe, except for PST, had weak to moderate positive correlations with the receptive language. This suggests that individuals who have faster and more efficient transmission of auditory-linguistic information to the cerebellum perform better on single-word comprehension tasks. This is reflected in our findings with the TP and the STG. The correlation of the MTG ROI may be a little less straight-forward. Some evidence has shown that the posterior MTG is involved in semantic control (Davey et al., 2016), while others have postulated that it could be involved in some form of integration between lobes (Turken & Dronkers, 2011). In addition, IFG also showed weak to moderate positive correlation with the task, which makes sense with its role in lexical and semantic processing (Dapretto & Bookheimer, 1999; Friederici, Opitz, & Von Cramon, 2000).

How selective are these findings? Our results show that there is some domain-specific connectivity to the cerebrum, even within large regions such as Crus I. Tracts from working memory ROIs in the cerebellum to language ROIs in the cerebrum were volumetrically smaller than the language-language tracts. This occurred even when the cerebellar ROIs from the different tasks were in the same lobule of the cerebellum. Some specificity was lost in the cortico-ponto-cerebellar tract (with fibers projecting to regions derived from the working memory task). Tracts seeded in IFG and projecting to working memory cerebellar regions Crus I, were not significantly different in volume from the tracts projecting to the language ROI (also located in Crus I). The longer tracts get, the less accurate they are, so this result may just be due to the fact that the regions of interest we had were located in the same lobule of the cerebellum. The specificity that these white matter tracts hold cannot be completely resolved without a more in depth knowledge of what these pathways truly look like in humans. In addition, our findings help localize what type of cerebellar damage should cause language comprehension deficits: lesions to right Crus I or Lobule IX, as well as to the polysynaptic fiber paths connecting these regions to the cerebrum. Individuals with lesions that preserve these regions are not expected to exhibit any language comprehension deficits.

### Limitations and Future Directions

First, the task set included in the HCP dataset is limited, which limited the cerebellar seed regions used in our tractography analysis. Future studies should test the limits of our findings by employing a range of language production and reception tasks involving lower-and higher-level processing, mapping the functional ROIs, and conducting tractography. Second, diffusion MRI tractography has been criticized for having a high rate of false positives (Maier-Hein et al., 2017; Thomas et al., 2014; Reveley et al., 2015). The best solution to this problem is to conduct histology-guided tractography analyses followed by replication in independent samples. Third, the relationship between structural connectivity, as described here, and functional connectivity, as reported in many prior studies (reviewed in Vias & Dick, 2017) in regards to the linguistic cerebellum needs to be investigated in a larger-scale study. Last, like all neuroimaging analyses, results can depend on various factors including the selection of ROIs, waypoints, thresholding, and software choices, etc. Every analytical decision made in this study was based on prior work exploring similar pathways (Metoki et al., 2021) with a small number of updates based on our understanding of best practices in diffusion imaging. However we are cognizant that diffusion imaging methods are relatively young, and standards are in flux. We hope that future investigators attempt replications of our work and that our findings are robust to different choices made in analytical parameters.

Future directions include the investigation of the specificity of the mentalizing and language cerebellar pathways. While it was shown that there was little functional overlap in the cerebellum between the language and working memory tasks, this was not the same for the language and mentalizing tasks. In order to further identify whether the language-specific pathways are truly language-specific, further research must be done between the language and mentalizing tasks in regards to diffusion tractography.

## Conclusions

This neuroimaging study investigated the structural connectivity between language cerebellar and cerebral areas. We found projections from the cerebellum to language regions of the frontal lobe. There was also evidence of input to the cerebellum from other regions of the cerebrum, such as the angular gyrus and the temporal pole portion of the superior temporal sulcus. These findings suggest that regions of the posterior cerebellum play a key role in language comprehension. We also found that these white matter tracts were, at least in part, specific to language-sensitive areas of the cerebellum and cerebrum.

## Acknowledgements

We would like to thank Angela Piecyk, Marah Dormuth, Sarah Johnson, Giovanna Arantes De Oliveira Campos, and Jason Konadu for their work as research assistants on this project. We would also like to thank Huiling Peng for assistance with tractography. Last, we would like to thank three anonymous reviewers for their helpful commentary.

