## Supplement for "Language and the cerebellum: structural connectivity to the eloquent brain"

**Supplementary Material**

Human Connectome Project protocol

The HCP protocol includes acquisition of structural MRI (0.7mm isotropic voxels), resting-state fMRI (rfMRI) (2mm isotropic voxels, four runs of 1200 volumes, TR = 720mm, 14min, 33sec per run), tfMRI (2mm isotropic voxels, TR = 720ms, tasks: working memory, gambling, motor, language processing, relational processing, social cognition, emotion processing), and dMRI (1.25mm isotropic voxels, b = 1000, 2000, and 3000s/mm2); 90 diffusion-weighting directions acquired with right-to-left and left-to-right phase encoding for each shell in a customized Siemens 3T Connectome Skyra scanner. Participants also received extensive behavioral testing (Barch et al. 2013).

In-scanner and out-of-scanner tasks

Participants were assessed in seven major domains in the scanner: 1) social cognition (mentalizing); 2) motor ability (visual, somatosensory, and motor systems); 3) gambling; 4) working memory/cognitive control systems and category specific representations; 5) language (semantic and phonological processing); 6) relational processing; and 7) emotion processing (Barch et al. 2013). In Jobson et al., we used data from the motor, working memory, and language processing tasks. The motor task was an adaptation of the task developed by Buckner and colleagues (2011; see also Yeo et al. 2011) and included finger, toe, and tongue movements. The working memory paradigm was a 2-back task with blocks of trials consisting of images of faces, places, tools, and body parts. The language processing task was developed by Binder et al. (2011) and consisted of brief auditory adapted stories from Aesop’s fables followed by a two-alternative forced-choice question asking participants about the topic of the story.

Outside the scanner, several tasks were administered, including a measure of receptive vocabulary, which refers to the comprehension of individual words. Participants listened to an audio recording of a word and four images on a computer screen. The objective of the task was to select the picture that matched the word. The task was administered so that each subsequent stimulus presentation was determined by how well the subject did on the last trial (Elam, 2021, Instrument: Language/Vocabulary Comprehension). This measure is often used as a developmental tool to measure receptive language ability, hence the adaptive format of the task (Gershon et al., 2013).

Image Preprocessing

The imaging data used in this article were the “minimally preprocessed” included in the WU-Minn HCP Consortium S900 Release (WU-Minn HCP Consortium 2015). The dMRI data preprocessing included echo planar imaging (EPI) distortion, eddy-current-induced distortion and subject motion correction, gradient nonlinearity correction, normalization of the b0 image intensity across runs, and registering the mean b0 volume to a native T1 volume. The tfMRI data had undergone spatial artifact/distortion correction, cross-modal registration, and spatial normalization to MNI space. Additionally, we further processed the dMRI data with FSL’s BEDPOSTX multi-shell, ball and stick model (Sotiropoulos et al. 2016) to model white matter fiber orientations and crossing fibers, and removed motion artifacts from the tfMRI data using ICA-AROMA (Griffanti et al., 2014; Pruim et al., 2015). All fMRI data were spatially smoothed at 4mm.

Transformation Matrices

As our analyses were performed in the subjects’ native spaces, specific care was needed to be used in the transformation of these spheres. Although the diffusion data was already formatted in T1w space, in the creation of these transformation matrices we treated the diffusion data as though it was still in diffusion-weighted space, so as to take extra care to ensure the seamlessness of our transforms. To delineate the transformation parameters, a series of linear registrations were performed. First, subjects’ b0 images extracted from the eddy corrected diffusion data were registered to their respective T1-weighted anatomical images with six degrees of freedom, a correlation ratio cost function and trilinear interpolation, yielding a diffusion to structural space conversion matrix. Then, six degrees of freedom, a correlation cost ratio and trilinear interpolation registration parameters were used to warp subjects’ T1-weighted images to the 2 x 2 x 2mm^3^ MNI space brain template, creating a structural to standard space conversion file. Next, the inverse of the two aforementioned matrices was taken to produce a structural to diffusion space and a standard to structural space conversion matrix, respectively. Last, to acquire the diffusion to standard space transformation parameters, the structural to standard space and diffusion to structural space matrices were concatenated. Finally, the inverse of this file was taken to create the standard to diffusion space conversion matrix that was used to convert the spheres to subject space. These transformation matrices were used for all native-to-standard and standard-to-native transformations included in our analyses. All data transformed into native space was evaluated for anatomical accuracy, and to ensure that the transformation matrices imposed no error on the resulting native-space ROIs. Note that this portion of the pipeline is extremely important - if you do not create these transformation matrices for your data, you will likely have low streamline counts or be unable to reconstruct these pathways. Transformation between modalities can be tricky, and should be approached with care. You cannot directly transform between standard space and diffusion space, often because those two modalities share nothing in common. By using a T1w image as the bridge between diffusion and standard data, one can have a more accurate transformation.

**Supplementary Tables**

Table A. MNI coordinates and associated z-scores from Neurosynth for the eight regions of interest included in this study. Each coordinate was used as the center of a 6mm sphere that was then subsequently used in probabilistic tractography. Please note that the coordinates for PST do not reflect the peak activation for the ‘posterior superior’ map that was generated from Neurosynth. The putative location of “Wernicke's Area” is highly variable across the literature and there is some controversy as to whether this term should even be used (Tremblay & Dick, 2016). However, we adjusted the original coordinates to be more consistent with the literature (e.g. moved it more anterior than the original coordinates (72, 37, 43 with a z-score of 11.82)). We used Heschel’s gyrus to guide this change and ensured that the location of the coordinates still presented a high z-score in Neurosynth.

| **Language ROI Coordinates** | | | | |
| --- | --- | --- | --- | --- |
| **Cerebral ROIs** | **MNI Coordinates** | | | |
|  | **x** | **y** | **z** | **z-score** |
| *STG* | -58 | -14 | 0 | 17.15 |
| *MTG* | -60 | -42 | 0 | 13.17 |
| *ITG* | -50 | -54 | -8 | 12.06 |
| *ANG* | -42 | -64 | 40 | 10.73 |
| *TP* | -44 | 12 | -24 | 13.07 |
| *IFG* | -44 | 20 | 16 | 9.99 |
| *DLPFC* | -46 | 34 | 32 | 9.51 |
| *PST* | -50 | -40 | 8 | 11.12 |
| **Cerebellar Language ROIs** | | | | |
| **Cerebellar ROI** | **MNI Coordinates** | | | |
|  | **x** | **y** | **z** | **percent attributed to location** |
| *Crus I (Language)* | 32 | -78 | -34 | 82% |
| *Lobule IX (Language)* | 6 | -52 | -42 | 87% |
| *Right VI (Working Memory)* | 32 | -60 | -30 | 64% |
| *Crus I (Working Memory)* | 40 | -58 | -32 | 99% |
| *Right VIIb (Working Memory)* | 40 | -64 | -50 | 48% |
| **Excluded Working Memory Cerebellar ROIs** | | | | |
| *Crus I* | 40 | -58 | -32 | 99% |
| *Crus I* | 34 | -56 | -32 | 64% |
| *Crus I* | 36 | -50 | -34 | 74% |
| *Lobule VI* | 32 | -60 | -28 | 86% |

Table B. Exclusion masks used in probabilistic tractography. All cerebral lobes are created from the Harvard Oxford cortical atlas. The peduncles were created from Johns Hopkins University ICBM-DTI-81 white-matter labels atlas.

| **Lobe of Interest for Tractography** | **Regions Included in Exclusion Mask** | |
| --- | --- | --- |
|  | **CTC** | **CPC** |
| *Frontal Lobe (IFG, DLPFC)* | Occipital lobe, parietal lobe, temporal lobe, right cerebral hemisphere, left cerebellar hemisphere & middle cerebellar peduncle | Occipital lobe, parietal lobe, temporal lobe, right cerebral hemisphere, left cerebellar hemisphere & superior cerebellar peduncle |
| *Parietal Lobe (Angular Gyrus)* | Occipital lobe, frontal lobe, temporal lobe, right cerebral hemisphere, left cerebellar hemisphere & middle cerebellar peduncle | Occipital lobe, frontal lobe, temporal lobe, right cerebral hemisphere, left cerebellar hemisphere & superior cerebellar peduncle |
| *Temporal Lobe (PST, STG, MTG, ITG, TP)* | Occipital lobe, parietal lobe, frontal lobe, right cerebral hemisphere, left cerebellar hemisphere & middle cerebellar peduncle | Occipital lobe, parietal lobe, frontal lobe, right cerebral hemisphere, left cerebellar hemisphere & superior cerebellar peduncle |

Table C. All reported Wilcoxon signed rank tests. CTC = cerebello-thalamo-cortical pathway; CPC = cortico-ponto-cerebellar pathway; IFG = inferior frontal gyrus/Broca’s; DLPFC = dorsomedial prefrontal cortex; ANG = angular gyrus; PST = posterior superior temporal; STG = superior temporal gyrus; MTG = middle temporal gyrus; ITG = inferior temporal gyrus; TP = temporal pole. Which tract was being assessed and which seed/target was being used (either Crus I or Lobule IX) is designated in parentheses after the ROIs being used for pairwise comparison. For example, when comparing two tracts that are seeded in Crus I and project to IFG and DLPFC, it will be labeled as: IFG & DLPFC (CTC - Crus I). All statistics were completed in RStudio. Adjusted p-values were also computed in R, with a total of 318 individual statistical tests that were completed for this paper and corrected using false detection rate (FDR). Note that for some comparisons, the rank assignment of the comparison was 0. Upon further investigation of the data, it became obvious that the reason for this was simply that one group had higher values than the other group for each data point. This results in a 0 rank sum, as there are either no positive or negative ranks to the data, making the average of the reported rank 0. R still reports a p-value in this case, which is shown below. Adjusted p-values that are significant at <0.001 are bolded. Adjusted r-values (effect size) which indicate a large effect at >0.5 are also in bold.

| **Pairwise Comparisons by ROI** | **Wilcoxon Signed Rank Test** | | | | |
| --- | --- | --- | --- | --- | --- |
|  | **n** | **V** | **p-value** | **adjusted p-value** | **adjusted r-value** |
| *ANG & IFG (CTC - Crus I)* | 100 | 0 | < 0.001 | **<0.001** | **0.737** |
| *ANG & DLPFC (CTC - Crus I)* | 100 | 0 | < 0.001 | **<0.001** | **0.737** |
| *ANG & ITG (CTC - Crus I)* | 100 | 2493 | 0.914 | 0.929 | 0.008 |
| *ANG & MTG (CTC - Crus I)* | 100 | 2381 | 0.662 | 0.731 | 0.030 |
| *ANG & STG (CTC - Crus I)* | 100 | 2462 | 0.83 | 0.823 | 0.015 |
| *ANG & TP (CTC - Crus I)* | 100 | 2649 | 0.671 | 0.759 | 0.027 |
| *ANG & PST (CTC - Crus I)* | 100 | 2316 | 0.473 | 0.643 | 0.040 |
| *IFG & DLPFC (CTC - Crus I)* | 100 | 699 | <0.001 | **<0.001** | 0.484 |
| *DLPFC & ITG (CTC - Crus I)* | 100 | 5050 | < 0.001 | **<0.001** | **0.729** |
| *DLPFC & MTG (CTC - Crus I)* | 100 | 5050 | < 0.001 | **< 0.001** | **0.729** |
| *DLPFC & STG (CTC - Crus I)* | 100 | 5050 | < 0.001 | **< 0.001** | **0.729** |
| *DLPFC & TP (CTC - Crus I)* | 100 | 5050 | < 0.001 | **< 0.001** | **0.729** |
| *DLPFC & PST (CTC - Crus I)* | 100 | 5050 | < 0.001 | **< 0.001** | **0.729** |
| *IFG & ITG (CTC - Crus I)* | 100 | 5050 | < 0.001 | **< 0.001** | **0.729** |
| *IFG & MTG (CTC - Crus I)* | 100 | 5050 | < 0.001 | **< 0.001** | **0.729** |
| *IFG & STG (CTC - Crus I)* | 100 | 5050 | < 0.001 | **<0.001** | **0.729** |
| *IFG & TP (CTC - Crus I)* | 100 | 5050 | < 0.001 | **<0.001** | **0.729** |
| *IFG & PST (CTC - Crus I)* | 100 | 5050 | < 0.001 | **<0.001** | **0.729** |
| *ITG & MTG (CTC - Crus I)* | 100 | 1336 | 0.030 | 0.073 | 0.156 |
| *ITG & STG (CTC - Crus I)* | 100 | 1857 | 0.445 | 0.621 | 0.043 |
| *ITG & TP (CTC -Crus I)* | 100 | 2705 | 0.075 | 0.153 | 0.124 |
| *ITG & PST (CTC - Crus I)* | 100 | 880 | < 0.001 | **<0.001** | 0.358 |
| *MTG & STG (CTC - Crus I)* | 100 | 2044 | 0.249 | 0.398 | 0.074 |
| *MTG & TP (CTC - Crus I)* | 100 | 3174 | < 0.001 | **<0.001** | 0.283 |
| *MTG & PST (CTC - Crus I)* | 100 | 1139 | 0.014 | 0.037 | 0.182 |
| *STG & TP (CTC - Crus I)* | 100 | 2852 | 0.020 | 0.037 | 0.170 |
| *STG & PST (CTC - Crus I)* | 100 | 1059 | 0.001 | 0.003 | 0.260 |
| *TP & PST (CTC - Crus I)* | 100 | 997 | < 0.001 | **< 0.001** | 0.383 |
| *ANG & IFG (CTC - Lobule IX)* | 100 | 222 | < 0.001 | **< 0.001** | **0.670** |
| *ANG & DLPFC (CTC - Lobule IX)* | 100 | 2159 | 0.209 | 0.352 | 0.081 |
| *ANG & ITG (CTC - Lobule IX)* | 100 | 4933 | < 0.001 | **< 0.001** | **0.700** |
| *ANG & MTG (CTC - Lobule IX)* | 100 | 4711 | < 0.001 | **< 0.001** | **0.636** |
| *ANG & STG (CTC - Lobule IX)* | 100 | 4447 | < 0.001 | **< 0.001** | **0.557** |
| *ANG & TP (CTC - Lobule IX)* | 100 | 2019 | 0.082 | 0.165 | 0.121 |
| *ANG & PST (CTC - Lobule IX)* | 100 | 4963 | < 0.001 | **< 0.001** | **0.708** |
| *IFG & DLPFC (CTC - Lobule IX)* | 100 | 5044 | < 0.001 | **<0.001** | **0.729** |
| *IFG & ITG (CTC - Lobule IX)* | 100 | 5050 | < 0.001 | **< 0.001** | **0.729** |
| *IFG & MTG (CTC - Lobule IX)* | 100 | 5050 | < 0.001 | **< 0.001** | **0.729** |
| *IFG & STG (CTC - Lobule IX)* | 100 | 5042 | < 0.001 | **< 0.001** | **0.729** |
| *IFG & TP (CTC - Lobule IX)* | 100 | 4901 | < 0.001 | **< 0.001** | **0.692** |
| *IFG & PST (CTC - Lobule IX)* | 100 | 5049 | < 0.001 | **< 0.001** | **0.729** |
| *DLPFC & ITG (CTC - Lobule IX)* | 100 | 4754 | < 0.001 | **< 0.001** | **0.648** |
| *DLPFC & MTG (CTC - Lobule IX)* | 100 | 4459 | < 0.001 | **< 0.001** | **0.560** |
| *DLPFC & STG (CTC - Lobule IX)* | 100 | 4160 | < 0.001 | **< 0.001** | 0.470 |
| *DLPFC & TP (CTC - Lobule IX)* | 100 | 2408 | 0.689 | 0.774 | 0.025 |
| *DLPFC & PST (CTC - Lobule IX)* | 100 | 4732 | < 0.001 | **< 0.001** | **0.642** |
| *ITG & MTG (CTC - Lobule IX)* | 100 | 1281 | < 0.001 | **< 0.001** | 0.349 |
| *ITG & STG (CTC - Lobule IX)* | 100 | 777 | 0.723 | **< 0.001** | **0.504** |
| *ITG & TP (CTC - Lobule IX)* | 100 | 9 | < 0.001 | **< 0.001** | **0.729** |
| *ITG & PST (CTC - Lobule IX)* | 100 | 2860 | 0.25 | 0.398 | 0.074 |
| *MTG & STG (CTC - Lobule IX)* | 100 | 1563 | 0.001 | 0.003 | 0.261 |
| *MTG & TP (CTC - Lobule IX)* | 100 | 16 | < 0.001 | **< 0.001** | **0.728** |
| *MTG & PST (CTC - Lobule IX)* | 100 | 4250 | < 0.001 | **< 0.001** | **0.527** |
| *STG & TP (CTC - Lobule IX)* | 100 | 18 | < 0.001 | **< 0.001** | **0.728** |
| *STG & PST (CTC - Lobule IX)* | 100 | 4616 | < 0.001 | **< 0.001** | **0.608** |
| *TP & PST (CTC - Lobule IX)* | 100 | 5029 | < 0.001 | **< 0.001** | **0.727** |
| *ANG & IFG (CPC - Crus I)* | 100 | 208 | < 0.001 | **<0.001** | **0.674** |
| *ANG & DLPFC (CPC - Crus I)* | 100 | 2148 | 0.195 | 0.330 | 0.085 |
| *ANG & ITG (CPC - Crus I)* | 100 | 4931 | < 0.001 | **<0.001** | **0.700** |
| *ANG & MTG (CPC - Crus I)* | 100 | 4708 | < 0.001 | **<0.001** | **0.635** |
| *ANG & STG (CPC - Crus I)* | 100 | 4428 | < 0.001 | **<0.001** | **0.551** |
| *ANG & TP (CPC - Crus I)* | 100 | 1998 | 0.070 | 0.147 | 0.126 |
| *ANG & PST (CPC - Crus I)* | 100 | 4958 | < 0.001 | **< 0.001** | **0.707** |
| *IFG & DLPFC (CPC - Crus I)* | 100 | 5045 | < 0.001 | **< 0.001** | **0.729** |
| *IFG & ITG (CPC - Crus I)* | 100 | 5050 | < 0.001 | **< 0.001** | **0.729** |
| *IFG & MTG (CPC - Crus I)* | 100 | 5050 | < 0.001 | **< 0.001** | **0.729** |
| *IFG & STG (CPC - Crus I)* | 100 | 5042 | < 0.001 | **< 0.001** | **0.729** |
| *IFG & TP (CPC - Crus I)* | 100 | 4914 | < 0.001 | **< 0.001** | **0.696** |
| *IFG & PST (CPC - Crus I)* | 100 | 5049 | < 0.001 | **< 0.001** | **0.729** |
| *DLPFC & ITG (CPC - Crus I)* | 100 | 4753 | < 0.001 | **< 0.001** | **0.648** |
| *DLPFC & MTG (CPC - Crus I)* | 100 | 4468 | < 0.001 | **<0.001** | **0.563** |
| *DLPFC & STG (CPC - Crus I)* | 100 | 4139 | < 0.001 | **<0.001** | 0.463 |
| *DLPFC & TP (CPC - Crus I)* | 100 | 2393 | 0.651 | 0.742 | 0.029 |
| *DLPFC & PST (CPC - Crus I)* | 100 | 4734 | < 0.001 | **< 0.001** | **0.643** |
| *ITG & MTG (CPC - Crus I)* | 100 | 1284 | < 0.001 | **< 0.001** | 0.348 |
| *ITG & STG (CPC - Crus I)* | 100 | 769 | < 0.001 | **< 0.001** | **0.507** |
| *ITG & TP (CPC - Crus I)* | 100 | 9 | < 0.001 | **<0.001** | **0.729** |
| *ITG & PST (CPC - Crus I)* | 100 | 2851 | 0.263 | 0.414 | 0.071 |
| *MTG & STG (CPC - Crus I)* | 100 | 1539 | 0.001 | 0.002 | 0.269 |
| *MTG & TP (CPC - Crus I)* | 100 | 17 | < 0.001 | **<0.001** | **0.728** |
| *MTG & PST (CPC - Crus I)* | 100 | 4352 | < 0.001 | **<0.001** | **0.609** |
| *STG & TP (CPC - Crus I)* | 100 | 18 | < 0.001 | **< 0.001** | **0.728** |
| *STG & PST (CPC - Crus I)* | 100 | 4622 | < 0.001 | **< 0.001** | **0.609** |
| *TP & PST (CPC - Crus I)* | 100 | 5025 | < 0.001 | **< 0.001** | **0.727** |
| *ANG & IFG (CPC - Lobule IX)* | 100 | 213 | < 0.001 | **< 0.001** | **0.673** |
| *ANG & DLPFC (CPC - Lobule IX)* | 100 | 2189 | 0.249 | 0.398 | 0.074 |
| *ANG & ITG (CPC - Lobule IX)* | 100 | 4922 | < 0.001 | **<0.001** | **0.697** |
| *ANG & MTG (CPC - Lobule IX)* | 100 | 4712 | < 0.001 | **< 0.001** | **0.636** |
| *ANG & STG (CPC - Lobule IX)* | 100 | 4453 | < 0.001 | **< 0.001** | **0.558** |
| *ANG & TP (CPC - Lobule IX)* | 100 | 2065 | 0.114 | 0.208 | 0.110 |
| *ANG & PST (CPC - Lobule IX)* | 100 | 4960 | < 0.001 | **< 0.001** | **0.708** |
| *IFG & DLPFC (CPC - Lobule IX)* | 100 | 5049 | < 0.001 | **< 0.001** | **0.729** |
| *IFG & ITG (CPC - Lobule IX)* | 100 | 5050 | < 0.001 | **< 0.001** | **0.729** |
| *IFG & MTG (CPC - Lobule IX)* | 100 | 5050 | < 0.001 | **< 0.001** | **0.729** |
| *IFG & STG (CPC - Lobule IX)* | 100 | 5042 | < 0.001 | **< 0.001** | **0.729** |
| *IFG & TP (CPC - Lobule IX)* | 100 | 4922 | < 0.001 | **< 0.001** | **0.697** |
| *IFG & PST (CPC - Lobule IX)* | 100 | 5049 | < 0.001 | **< 0.001** | **0.737** |
| *DLPFC & ITG (CPC - Lobule IXI)* | 100 | 4753 | < 0.001 | **< 0.001** | **0.648** |
| *DLPFC & MTG (CPC - Lobule IX)* | 100 | 4442 | < 0.001 | **< 0.001** | **0.555** |
| *DLPFC & STG (CPC - Lobule IX)* | 100 | 4146 | < 0.001 | **< 0.001** | 0.465 |
| *DLPFC & TP (CPC - Lobule IX)* | 100 | 2426 | 0.735 | 0.795 | 0.023 |
| *DLPFC & PST (CPC - Lobule IX)* | 100 | 4723 | < 0.001 | **< 0.001** | **0.640** |
| *ITG & MTG (CPC - Lobule IX)* | 100 | 1290 | < 0.001 | **< 0.001** | 0.347 |
| *ITG & STG (CPC - Lobule IX)* | 100 | 831 | < 0.001 | **< 0.001** | 0.488 |
| *ITG & TP (CPC - Lobule IX)* | 100 | 10 | < 0.001 | **< 0.001** | **0.729** |
| *ITG & PST (CPC - Lobule IX)* | 100 | 2866 | 0.242 | 0.395 | 0.074 |
| *MTG & STG (CPC - Lobule IX)* | 100 | 1621 | 0.002 | 0.006 | 0.241 |
| *MTG & TP (CPC - Lobule IX)* | 100 | 19 | < 0.001 | **< 0.001** | **0.728** |
| *MTG & PST (CPC - Lobule IX)* | 100 | 4403 | < 0.001 | **< 0.001** | 0.544 |
| *STG & TP (CPC - Lobule IX)* | 100 | 10 | < 0.001 | **< 0.001** | **0.729** |
| *STG & PST (CPC - Lobule IX)* | 100 | 4594 | < 0.001 | **< 0.001** | **0.601** |
| *TP & PST (CPC - Lobule IX)* | 100 | 5027 | < 0.001 | **< 0.001** | **0.727** |
| *Crus I (language) & Lobule VIIb (working memory) (CTC - IFG)* | 100 | 3988 | <0.001 | **<0.001** | 0.417 |
| *Crus I (language) & Crus I (working memory) (CTC - IFG)* | 100 | 4913 | < 0.001 | < 0.001 | **0.696** |
| *Crus I (language) & Lobule IV (working memory) (CTC - IFG)* | 100 | 4988 | <0.001 | **<0.001** | **0.716** |
| *Lobule IX (language) & Lobule VIIb (working memory) (CTC - IFG)* | 100 | 4116 | <0.001 | **<0.001** | 0.457 |
| *Lobule IX (language) & Crus I (working memory) (CTC - IFG)* | 100 | 4938 | <0.001 | **<0.001** | **0.701** |
| *Lobule IX (language) & Lobule IV (working memory) (CTC - IFG)* | 100 | 4955 | <0.001 | **<0.001** | **0.706** |
| *Crus I (language) & Lobule VIIb (working memory) (CPC - IFG)* | 100 | 3698 | <0.001 | **<0.001** | 0.327 |
| *Crus I (language) & Crus I (working memory) (CPC - IFG)* | 100 | 2725 | 0.493 | 0.667 | 0.037 |
| *Crus I (language) & Lobule IV (working memory) (CPC - IFG)* | 100 | 3770 | <0.001 | **<0.001** | 0.349 |
| *Lobule IX (language) & Lobule VIIb (working memory) (CPC - IFG)* | 100 | 4743 | <0.001 | **<0.001** | **0.645** |
| *Lobule IX (language) & Crus I (working memory) (CPC - IFG)* | 100 | 3932 | <0.001 | **<0.001** | 0.399 |
| *Lobule IX (language) & Lobule IV (working memory) (CPC - IFG)* | 100 | 4559 | <0.001 | **<0.001** | **0.591** |
